# Global hypoperfusion leads to a mismatch in oxygen delivery and consumption in the cerebral watershed area

**DOI:** 10.1101/2025.06.16.659854

**Authors:** Baoqiang Li, Hewei Cao, Hajime Takase, Srinivasa Rao Allu, Yimeng Wu, Buyin Fu, Sergei A. Vinogradov, Ken Arai, Eng H. Lo, Cenk Ayata, Sava Sakadžić

## Abstract

Despite the pivotal role of pial collaterals in maintaining cerebral blood flow during focal brain ischemia, it is largely unexplored how the microvascular blood flow and oxygenation in the watershed “pial-collateral territory” differ from those in the territory supplied by the major arteries during chronic global hypoperfusion. To answer this question, we applied 2-photon microscopy and Doppler optical coherence tomography to investigate the changes in cerebral microvascular blood flow and partial pressure of oxygen (PO_2_), induced by bilateral common carotid artery stenosis (BCAS). The measurements were performed in the somatosensory cortex that is supplied by the middle cerebral artery (MCA), and in the adjacent watershed area in the awake, head-restrained C57BL/6 mice, via the chronic cranial window. The results showed that the BCAS induced a larger decrease in capillary red blood cell (RBC) flux in the watershed area than in the MCA territory, especially in the subcortical white matter. Besides, PO_2_ in the pial collaterals was significantly lower than that in the upstream MCA segments under control conditions. However, the PO_2_ changes in the arteries and veins under global hypoperfusion displayed different trends in the two interrogated regions, resulting in a significant increase in oxygen extraction fraction in the watershed area. These findings suggest a mismatch between oxygen supply and demand in the watershed area due to global hypoperfusion and increased subcortical white matter vulnerability. We have also observed dilation of the pial collaterals after BCAS, which might suggest a compensatory mechanism to improve the blood flow in the watershed under hypoperfusion.

## INTRODUCTION

Collateral circulation has shown its potential in optimizing treatment and management strategies for cerebral ischemia, and attracted increasing attention in stroke research.^1–3^ Collaterals refer to the alternative vascular pathways to deliver oxygen and nutrients to the hypoperfused tissue when the principal routes are impaired, for instance, in cases of large-vessel occlusion and high-grade stenosis.^4–7^ Intracranial collaterals mainly consist of the Circle of Willis (CoW) and the leptomeningeal anastomosis vessels (a.k.a., pial collaterals), which are regarded as the primary and secondary collaterals, respectively.^8^ CoW is a ring of arterial segments, providing anastomotic connections between the anterior and posterior circulations.^9^ Pial collaterals refer to the small arterial segments connecting the terminal branches of the major arteries at the pial surface.^10,11^ Although CoW is capable of providing large-scale diversion of blood flow, the pial collaterals ultimately deliver the oxygenated blood to the hypoperfused tissue on a much finer scale, demonstrating their critical role, although small in size, in sustaining cerebral perfusion during acute ischemia or chronic hypoperfusion.^12–15^

The spatiotemporal pattern of pial-collateral circulation has been increasingly studied in animal models. Investigators observed collateral recruitment during ischemic stroke and chronic hypoperfusion, finding that pial-collateral circulation was strongly correlated with brain injury severity and behavioral deficits,^16–19^ and that abundant pial collaterals led to smaller brain infarcts and better treatment outcomes.^20,21^ On the other hand, the associated border-zone cortical tissue area (a.k.a., watershed), which typically situates underneath the pial-collateral networks, is susceptible to hemodynamic insufficiency. Clinical studies showed that carotid artery stenosis limited cerebral perfusion downstream along the arterial tree, causing ischemic lesions in the watershed area (a.k.a., watershed infarcts), especially in the deep subcortical white matter.^22,23^ This emphasizes the importance of better understanding the regulations of blood flow and oxygen delivery in the watershed area. However, how microvascular blood flow and oxygen are distributed in the watershed tissue area under normal and physiologically disturbed conditions as well as how these distributions differ from those in the territory supplied by the major arteries remain largely unknown.

In the present work, we applied a set of advanced imaging techniques to answer these questions. Specifically, we used 2-photon microscopy (2PM) to measure intravascular partial pressure of oxygen (PO_2_), capillary red-blood-cell (RBC) flux, and to acquire microvascular angiograms. In addition, we used Doppler optical coherence tomography (Doppler-OCT) to measure absolute cerebral blood flow. These measurements were performed in the somatosensory cortex that is supplied by the middle cerebral artery (MCA) and for the very first time in the adjacent watershed area, in the awake, head-restrained C57BL/6 mice. The imaging experiments were conducted under control conditions (i.e., normal physiology) and global hypoperfusion that was induced by bilateral common carotid artery stenosis (BCAS). Several novel observations were made with these multifaceted measurements. Briefly, we found that the BCAS induced differentially smaller RBC flux in the watershed area, especially in the subcortical white matter. The baseline PO_2_ in the pial collaterals was significantly lower than that in the MCA segments. However, the changes in the arterial and venous PO_2_ in the two interrogated regions exhibited different trends under global hypoperfusion, giving rise to a significant increase in oxygen extraction fraction (OEF) in the watershed area. We have also observed dilation of the pial collaterals, which might play a compensatory role in ameliorating a large mismatch between oxygen delivery and consumption in the watershed. This work reports regional differences in the distributions of cerebral microvascular blood flow and oxygenation in the watershed area vs. in the MCA territory. It provides novel insights into how they are redistributed during chronic global hypoperfusion, an evidence of O_2_ delivery-consumption mismatch, higher vulnerability of watershed and particularly subcortical white matter in the watershed to hypoperfusion, and potentially a novel role of pial collaterals in mitigating the effects of hypoperfusion in the watershed.

## MATERIAL AND METHODS

### Animal preparation

We used n=8 C57BL/6 mice (female, 3-5-month old, 20-30-gram body weight; Charles River Laboratories) in this work. For each mouse, a cover-glass-sealed, round shape (3 mm in diameter), chronic cranial window was implanted, centering approximately over the E1 whisker barrel in the left hemisphere (AP: -1.5 mm, ML: -3.0 mm relative to bregma), following the published procedures.^24–26^ A custom-made aluminum head- post was glued to the skull over the right hemisphere. During imaging experiments, the head-post was attached to a home-built cradle, allowing head immobilization.^27^ After cranial surgery, the mice were given 5 days to recover. Subsequently, the mice were gradually habituated, during 2-3 weeks, to increasingly extended periods of head-restraint in the cradle (e.g., from 10 minutes to 2 hours). The optical measurements were first performed under control conditions. Three weeks after the control measurements were completed, the BCAS surgery was conducted in each mouse following the previously established protocol.^28^ The same mice were imaged again 7 days after the BCAS surgery for comparisons.

All the surgical and experimental procedures were approved by the Massachusetts General Hospital Institutional Animal Care and Use Committee, in accordance with the National Institutes of Health’s Guide for the Care and Use of Laboratory Animals.

### 2PM setup

A home-developed 2-photon laser scanning microscope was used in this work. Briefly, the system was equipped with a commercial laser system (680–1,300 nm tuning range, ∼120 fs pulse width, 80 MHz pulse repetition rate; InSight DeepSee, Spectra-Physics). The laser power was controlled by an electro-optic modulator (EOM) (ConOptics). The laser beam was scanned in the X-Y plane by a pair of galvanometer mirrors (6215H, Cambridge Technology). A water-immersion 20X objective lens (1.0 NA; XLUMPLFLN20XW, Olympus) and an air-spaced 4X objective lens (0.28 NA; XLFLUOR4X/340, Olympus) were used for imaging. A motorized stage (M-112.1DG, Physik Instrumente) was employed to translate the objective lens along the Z-axis to focus the laser beam at different focal planes. The detection path was comprised of a dichroic mirror (FF875-Di01-38.1×51.0, Semrock) followed by an infrared blocker (FF01-890/SP-50, Semrock), an emission filter (FF01-795/150-25 or FF01-709/167-25 for phosphorescence or fluorescence imaging, respectively, Semrock), and a photon-counting photomultiplier tube (PMT) (H10770PA-50, Hamamatsu). The PMT signal was input into a discriminator (C9744, Hamamatsu) and then sampled by a digital acquisition card (PCle-6537, National Instruments). During experiments, the water-immersion 20X objective lens was heated by an electric heater (TC-HLS-05, Bioscience Tools) to maintain the temperature of the water medium close to ∼37 °C. More details about the 2PM system can be found in our previous works.^29,30^

### Imaging of capillary RBC flux

To measure RBC flux, blood plasma was labeled with a near-infrared emitting fluorophore – Alexa680 conjugated with 70-kDa dextran molecules. The maximum 2P excitation wavelength (2P-λ_exc_) and the peak wavelength of the emission spectrum (λ_em_) of Alexa680 are 1,280 nm and 700 nm, respectively. The dextran-Alexa680 solution (0.1 mL at 5% W/V in PBS) was injected retro-orbitally into the bloodstream under brief isoflurane anesthesia (1.5-2%, during ∼2 minutes) ∼20 minutes before the imaging experiment. We used 2PM with the 20X objective lens for RBC-flux measurements. In each mouse, multiple regions of interest (ROIs) were selected, typically with one ROI chosen in the MCA territory and one or two ROI(s) in the adjacent watershed area, where the pial collaterals could be empirically identified (Fig. 1A). RBC flux was measured in the cortical gray matter at the depths of 0.4 mm (corresponding to layer IV) and 0.6 mm (layer V) under the cortical surface, as well as at 3-5 planes with 10-μm intervals in the subcortical white matter with an average imaging depth of ∼0.82 mm. The depths of the white matter boundaries were determined by an OCT-based method.^31^ At each imaging depth, a fluorescence-labelled angiogram was acquired to reveal the microvascular structure within a 0.5×0.5 mm^2^ field of view (FOV). Guided by the angiographic image (Fig. 1B), we manually selected the measurement locations in most capillaries that could be visually inspected, with one measurement location in each capillary. Here, capillaries were identified empirically based on vascular caliber. While imaging, the laser beam was parked at each measurement location for 0.5 s. The emitted fluorescence was detected by the PMT operated in the photon-counting mode. The acquired fluorescence signal was binned using 250-μs-wide bins, yielding a 2,000-point (corresponding to a 0.5-s-long duration) fluorescence intensity time course (Fig. 1C). The amplitude of fluorescence intensity could be adjusted by the laser power, which was controlled by the EOM and kept constant for imaging in the same depth. Since the dextran-Alexa680 molecules were dissolved in the blood plasma and, therefore, did not label the RBCs, the fluorescence intensity changes encoded the flowing of RBCs and blood plasma, as indicated by the “valleys” and “peaks” in the representative fluorescence intensity time courses in Fig. 1C. The fluorescence intensity time courses were segmented with a previously-established binary thresholding approach.^31^ The segmentation was evaluated by the coefficient of determination (R^2^) between the experimental and fitted time courses, and the time courses with R^2^<0.5 were rejected. This criterion ensures that the selected time courses are recorded from the vessels with single-file RBC flowing profiles, which are mostly capillaries. Finally, RBC flux was explicitly calculated by counting the number of RBC passages normalized by the acquisition time.

**Fig. 1.**
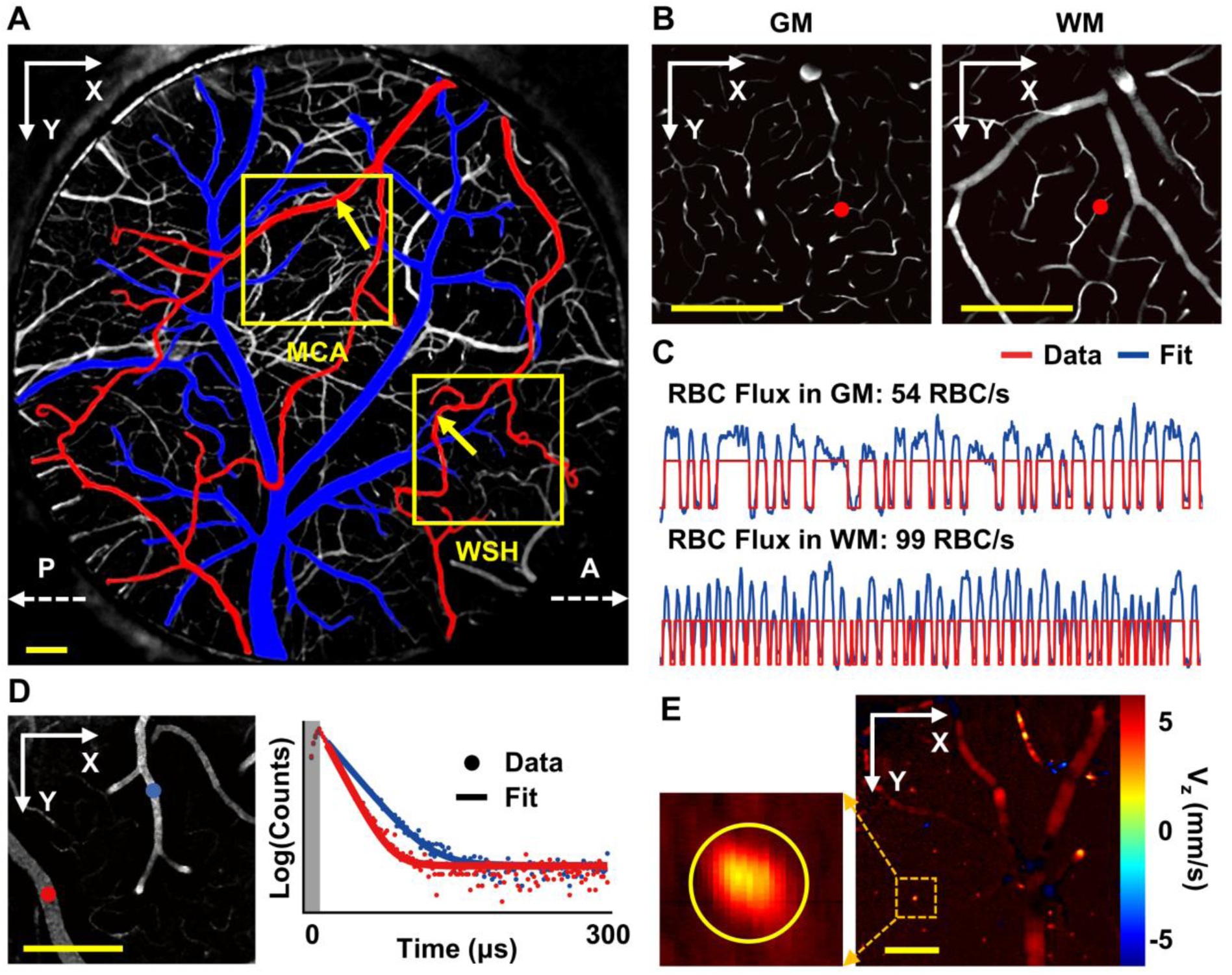
Imaging methods. **A.** Maximum intensity projection (depth range of 0-100 μm) of the Alexa680-labeled angiogram (3.0×3.0 mm^2^ FOV) acquired using 2PM with the 4X objective lens in a representative animal. The anatomical orientation of this image is indicated by the dashed white arrows (A: Anterior; P: Posterior). The primary arteries (including pial collaterals) and veins at the pial surface are color-coded in red and blue, respectively. The two yellow squares enclose the representative ROIs for imaging in the MCA territory and watershed area (WSH), with the MCA and pial-collateral segments connecting the terminal branches of the anterior and middle cerebral arteries denoted by the yellow arrows. **B.** Angiographic images were acquired in the cortical gray matter (GM) at a depth of 0.6 mm and in the subcortical white matter (WM) at 1.05 mm. **C.** Two representative 0.5-s-long fluorescence intensity time courses were acquired in the GM and WM capillaries with the measurement locations denoted by the red dots in panel **B**. The flux values are calculated as 54 RBC/s and 99 RBC/s, respectively. **D.** Two representative phosphorescence decay curves (photon counts in logarithmic scale) for PO2 recording in an arteriole (in red; lifetime=12.3 μs, PO2=97 mmHg) and a venule (in blue; lifetime=19.4 μs, PO2=43 mmHg). **E.** An en-face Doppler- OCT image acquired at the cortical surface (right), and the zoomed-in view of a surfacing venule (left) enclosed by the yellow circle. Scale bars: 200 μm.

### PO_2_ imaging

We used an oxygen-sensitive phosphorescence probe – Oxyphor2P for PO_2_ imaging.^32^ The 2P-λ_exc_ and λ_em_ of Oxyphor2P are 950 nm and 757 nm, respectively. The Oxyphor2P solution (0.1 mL at 34 µM in PBS) was injected retro-orbitally into the bloodstream under brief isoflurane anesthesia (1.5-2%, during ∼2 minutes) ∼20 minutes before the imaging experiment.

We used 2PM with the 20X objective lens for PO_2_ imaging. The intravascular PO_2_ measurements were performed from the cortical surface down to 0.5 mm under the cortical surface (layer V) with 0.1-mm intervals, in the same ROIs as for RBC-flux imaging. At each imaging depth, a raster scan was performed, and a phosphorescence intensity image was acquired to reveal the microvascular structure within a 0.5×0.5 mm^2^ FOV (left side in Fig. 1D). We then manually selected the measurement locations in most of the vascular segments including arterioles, venules, and capillaries, with one measurement location in each segment. The arteriolar segments in the watershed area are mostly branched from the pial collaterals. For measuring the phosphorescence lifetime, the Oxyphor2P molecules were excited with a 10-µs-long laser excitation gated by the EOM, followed by a 290-µs-long recording of the emitted phosphorescence photons. The laser beam was parked at each location to repeat such 300-µs-long excitation/detection cycle for 2,000 times (corresponding to a 0.6-s-long duration) to obtain an average phosphorescence decay curve (right side in Fig. 1D). The phosphorescence lifetime was thus calculated by fitting the decay curve to a single-exponential decay function. Finally, the phosphorescence lifetime was converted into absolute PO_2_ based on the Stern-Volmer type calibration.^29,32^

### OCT imaging

A spectral-domain OCT system was employed in this study. The system was equipped with a dual- superluminescent-diode light source (central wavelength=1310 nm, bandwidth=170 nm; LS2000B, Thorlabs). The axial resolution, pixel size, and imaging range along the axial direction were experimentally estimated as 3.5 µm, 2.8 µm, and 1.6 mm in biological tissue, respectively. With a 10X objective lens (Plan Apo NIR, Mitutoyo), the transverse resolution was estimated as 3.5 µm. More details about this system are available in our previous works.^33–35^

In each mouse, the Doppler-OCT measurements were performed in a 1×1 mm^2^ FOV, with the scan steps of 0.26 μm and 1.95 μm along the fast (X) and slow (Y) scanning axes, respectively. The Doppler- OCT images were acquired in the ROIs that covered the ROIs selected for PO_2_ and RBC-flux imaging. With the Doppler-OCT images, flow speed along the Z-axis (V_z_; Fig. 1E) was computed with a Kasai autocorrelation algorithm.^36^ The imperfect telecentricity of the optical system would bring about an “offset” speed component along the Z-axis, which could be corrected using a previously-established method.^37,38^ Blood flow was then calculated by integrating the corrected V_z_ values over a manually drawn geometrical shape (e.g., the yellow circle in Fig. 1E) that fully encloses the cross-section of the selected vessel. In this work, only the flow data in venules were included into the analysis, because the measurements in arterioles might be affected by the aliasing effect caused by high flow speed. Twenty volumetric scans were repeated in each ROI for averaging.

### Calculation of SO_2_ and OEF

Oxygen saturation of hemoglobin (SO_2_) was calculated using the Hill equation with the parameters h=2.59 and P_50_=40.2 mmHg, which are specific for C57BL/6 mice.^39^ Here, h is the Hill coefficient, and P_50_ is the PO_2_ at which hemoglobin is half-saturated. Oxygen extraction fraction (OEF) is defined as (SO_2,A_– SO_2,V_)/SO_2,A_, where SO_2,A_ and SO_2,V_ represent the arterial and venous SO_2_, respectively.

### Angiogram and morphological analysis

We applied 2PM with the 20X objective lens to image the Alexa680-labeled microvasculature in a 0.7×0.7 mm^2^ FOV, with the scan steps of ∼1.37 μm in the X and Y directions. An angiogram stack was constructed by acquiring such two-dimensional image slices every 2 μm along the Z-axis from the cortical surface down to the subcortical white matter, in the brain regions that covered the ROIs selected for the PO_2_ and RBC- flux measurements. In a sub-group of the mice, the Alexa680-labeled angiograms were also acquired using 2PM with the 4X objective lens in a 3.0×3.0 mm^2^ FOV, covering the entire cranial window, with scan steps of ∼5.86 μm in the X and Y directions. Similarly, an angiogram stack was constructed by acquiring such two-dimensional image slices every 5 μm along the Z-axis from the cortical surface down to 100 μm under the cortical surface.

The high-resolution angiograms acquired with the 20X objective lens were used for morphological analysis. First, an angiographic image was segmented into a binary mask using a deep-learning-based algorithm.^40^ Then, a Laplacian optimization algorithm was used to generate a graph-based anatomical model of the microvasculature.^41^ Thus, morphological characteristics, such as vascular diameter, segment length, and segment density, could be calculated. Here, vascular diameter was calculated as the full width at half maximum of the profile along the radial direction of a vessel; segment length was calculated along the vessel centerline between the bifurcations at the two ends of a vessel segment; segment density was calculated as the number of vessel segments in a unit volume of tissue. This analysis included the micro- vessels with vascular diameter ≤10 µm (e.g., capillaries, and the smallest arterioles and venules) and the larger vessels with vascular diameter >10 µm, with the images acquired at the depth range of 100-700 μm under the cortical surface. The image data acquired at >700 μm were too noisy to be segmented accurately and, therefore, excluded from the analysis. In addition, the vascular diameter of the vessels at the pial surface (e.g., pial arteries, venues, and collaterals) was also calculated.

### Construction of the composite image

The composite images were constructed with the Alexa680-labeled angiograms acquired with the 20X objective lens and the experimental PO_2_ measurements. An angiographic image was first segmented into a binary mask using a deep-learning-based algorithm.^40^ Then, a Laplacian optimization algorithm was used to generate an anatomical graph of the microvasculature.^41^ With the graph, different vessel types (e.g., arteries, veins, and capillaries) were color-coded for visualization. Besides, PO_2_ data could be spatially co-registered to the graph. Here, the PO_2_ value obtained at the measurement location in a vascular segment was assigned to the whole segment. A composite PO_2_ image was thus generated by color-coding the PO_2_ values in the corresponding vascular segments.

### Statistical analysis

The study design and reporting of this work are in accordance with the ARRIVE guidelines. All data are presented as mean±STD, where applicable. Statistical comparisons were conducted with the Student’s t-test, and the P value ≤0.05 was considered statistically significant. Sample sizes were chosen to detect 30% difference between the mean values of each parameter (coefficient of variance=0.2, power=0.8, and α=0.05), and specified in the text and/or figure legends, where relevant. For each parameter, the relative change is defined as the ratio (in percentage) of the difference in the measured values between the control condition and the BCAS-induced hypoperfusion to that under the control condition, and the mean value was first calculated over the measurements performed in each mouse, and then across mice.

## RESULTS

### BCAS induced a significantly larger decrease in microvascular blood flow in the watershed area than in the MCA territory

We applied Doppler-OCT to investigate the changes in venous blood flow induced by BCAS. Fig. 2A shows that blood flow in the MCA territory decreased from 0.062±0.041 µL/min under the control conditions to 0.052±0.029 µL/min under the BCAS-induced global hypoperfusion, with a relative change of 11.8±26.1%. A trend of larger flow decrease was observed in the watershed area, i.e., from 0.043±0.023 µL/min under control condition to 0.035±0.015 µL/min under global hypoperfusion, with a relative change of 15.1±10.8%.

**Fig. 2.**
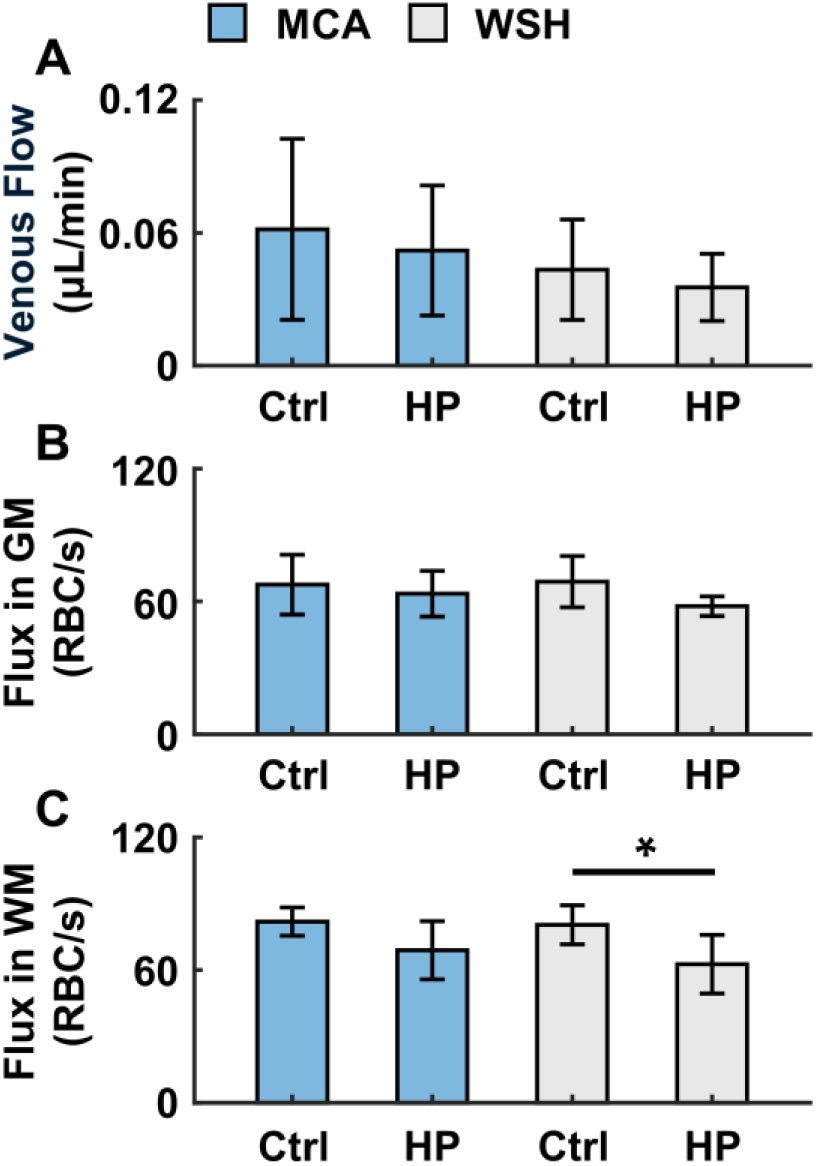
BCAS-induced decrease in microvascular blood flow. **A.** Absolute blood flow measured with Doppler-OCT in the ascending venules close to the cortical surface. The measurements under control condition (Ctrl) were performed in 43 vessels in the MCA territory and 75 vessels in the watershed area (WSH), in n=4 mice. The same vessels were selected for analysis under the BCAS-induced global hypoperfusion (HP). The mean flow value for each ROI (e.g., MCA or WSH) and under each condition (e.g., Ctrl or HP) was calculated first over the vessels selected in each mouse, and then across mice. **B, C.** Comparisons of RBC flux measured in the cortical gray matter (GM; panel **B**) and subcortical white matter (WM; panel **C**). The measurements in the GM were performed in 299 and 239 capillaries in the MCA territory under control condition and global hypoperfusion, respectively, as well as in 525 and 428 capillaries in the watershed area under control condition and global hypoperfusion, respectively. The measurements in the WM were performed in 282 and 154 capillaries in the MCA territory under control condition and global hypoperfusion, respectively, as well as in 265 and 381 capillaries in the watershed area under control condition and global hypoperfusion, respectively. The data presented in panels **B** and **C** were acquired in n=4 mice. In each mouse, RBC flux was measured at 0.4 mm (corresponding to layer IV) and 0.6 mm (layer V) in GM, as well as in WM with an average imaging depth of 0.82 mm. The measurements under the two conditions were performed approximately at the same depths and in the same ROIs. The mean flux for each ROI (e.g., MCA or WSH) and under each condition (e.g., Ctrl or HP) was calculated first over the capillaries selected in each mouse, and then across mice. Data are expressed as mean±STD. The asterisk symbol indicates statistical significance (Student’s t-test, P<0.05).

We further applied 2PM to investigate the BCAS-induced changes of RBC flux in the cortical gray matter (GM) and subcortical white matter (WM). Fig. 2B shows that RBC flux measured in the GM capillaries in the MCA territory was slightly decreased from 67.6±13.5 RBC/s under the control conditions to 63.5±10.3 RBC/s under global hypoperfusion, with a small relative change of 3.7±22.1%. A more pronounced RBC-flux decrease was observed in the watershed area, i.e., from 68.9±11.6 RBC/s under the control conditions to 57.8±4.4 RBC/s under global hypoperfusion, with a relative change of 15.0±8.8%. But no statistical significance was found (P=0.06). Shown in Fig. 2C, RBC flux measured in the WM capillaries in the MCA territory was decreased from 81.9±6.4 RBC/s under the control conditions to 68.9±13.2 RBC/s under global hypoperfusion, with a relative change of 15.1±20.4%. A significantly larger decrease was seen in the watershed area, i.e., from 80.4±8.8 RBC/s under the control conditions to 62.6±13.3 RBC/s under global hypoperfusion (P=0.046), exhibiting a large relative change (i.e., 22.6±11.0%).

### Oxygen extraction fraction in the watershed area was significantly increased under global hypoperfusion

We applied 2PM with Oxyphor2P to investigate the changes in cerebral microvascular oxygenation induced by BCAS. The composite images in Fig. 3A illustrate the PO_2_ distributions in the pial vessels in a representative animal. As shown, PO_2_ measured in the pial collaterals (WSH) is lower than that in the upstream MCA segments (MCA) under the control conditions. In the MCA territory, the changes of arterial and venous PO_2_ induced by global hypoperfusion were negligible. While, in the watershed area, noticeable trends of PO_2_ increase in the pial collaterals and PO_2_ decrease in the pial veins could be observed.

**Fig. 3.**
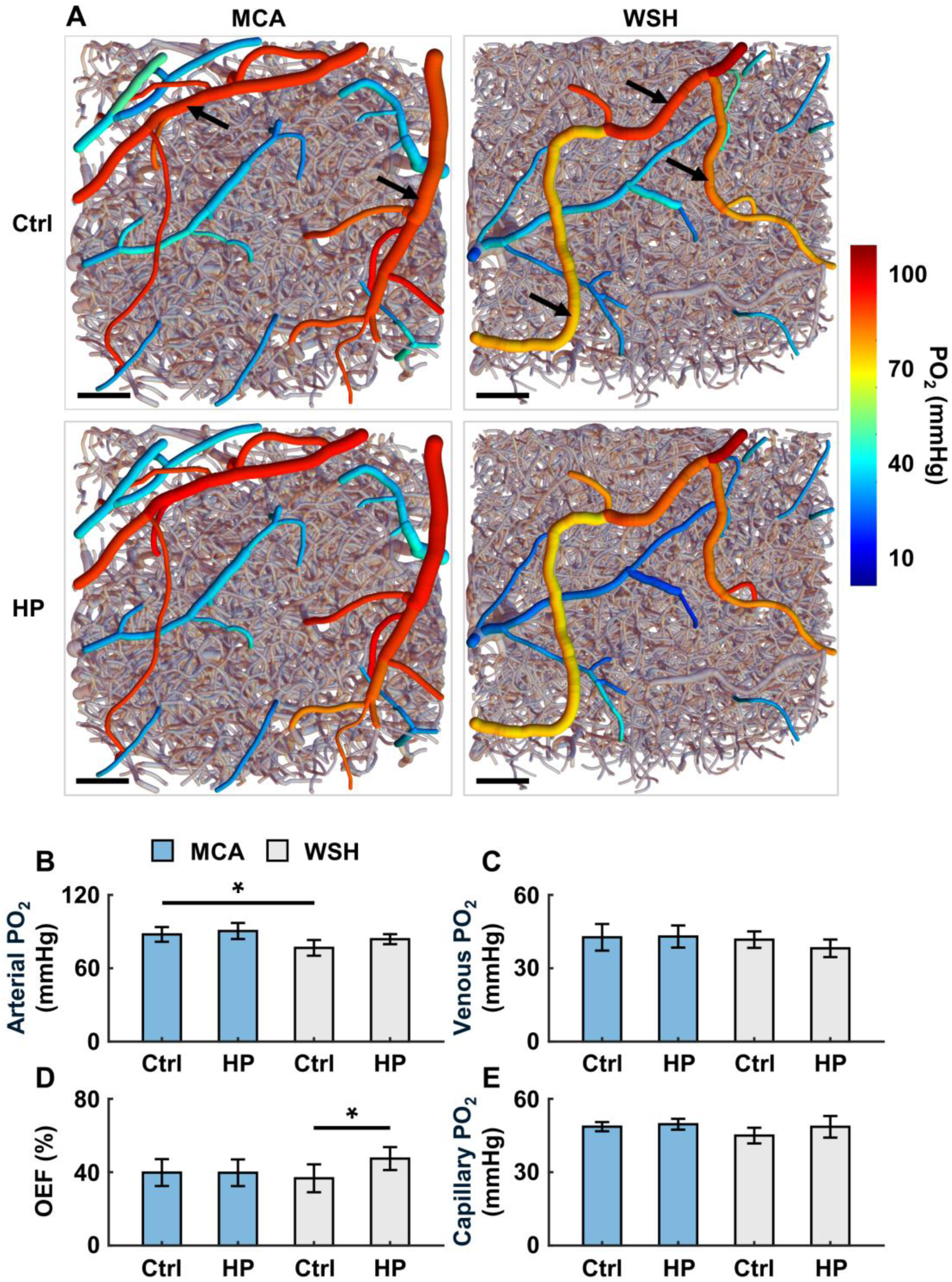
BCAS-induced changes of microvascular oxygenation. **A.** Composite images showing the PO2 distributions in the pial vessels under both conditions in a representative animal. The anatomical graphs, which illustrate the top view (X-Y plane) of the microvasculature from the cortical surface down to 600 μm, were generated from the angiograms acquired in the ROIs (e.g., MCA and WSH) as indicated by the yellow squares in Fig. 1A. The PO2 values obtained in the pial vessels (including arteries, veins, and pial collaterals) are color-coded. The vessels without PO2 data were set to be in gray color and half-transparent. In each ROI (e.g., MCA or WSH), the vascular graph was generated from the angiogram acquired under the control conditions, and that same graph was used as a template to co-register with the PO2 data acquired under the two different conditions separately. The black arrows in the MCA and WSH images under the control conditions denote representative MCA and pial-collateral segments, respectively. Scale bars: 100 μm. **B, C.** Comparisons of the arteriolar and venular PO2. The measurements were taken in the pial arterioles (**B**) and venules (**C**), as well as in the penetrating ones branched from the pial vasculature close to the cortical surface. The arteriolar segments selected in the watershed area mostly belong to the pial collaterals. The measurements under control condition were performed in 20 arterioles and 35 venules in the MCA territory, as well as in 34 arterioles and 44 venules in the watershed area, in n=5 mice. The same vessels were selected for measurements under global hypoperfusion. The mean PO2 value for each ROI (i.e., MCA or WSH) and under each condition (i.e., Ctrl or HP) was calculated first over the vessels selected in each mouse, and then across mice. **D.** Oxygen extraction fraction (OEF) that was calculated with the same dataset presented in panels **B** and **C**. **E.** Comparisons of capillary PO2 with the measurements performed at the depth from 100 μm down to 500 μm under the cortical surface, in n=5 mice. The measurements were performed in 582 and 494 capillaries in the MCA territory under control condition and global hypoperfusion, respectively, as well as in 1403 and 1158 capillaries in the watershed area under control condition and global hypoperfusion, respectively. In each mouse, the measurements under the two conditions were performed approximately at the same depths and in the same ROIs. The mean PO2 for each ROI (e.g., MCA or WSH) and under each condition (e.g., Ctrl or HP) was calculated first over the capillaries selected in each mouse, and then across mice. Data are expressed as mean±STD. The asterisk symbol indicates statistical significance (Student’s t-test, P<0.05).

The microvascular PO_2_ changes were further quantified. Shown in Fig. 3B, arterial PO_2_ in the MCA territory slightly increased from 87.7±6.0 mmHg under the control conditions to 90.5±6.6 mmHg under global hypoperfusion, with a relative change of 3.6±11.2%. While in the watershed area, arterial PO_2_, which was measured mostly in pial collaterals, increased from 76.7±6.4 mmHg under the control conditions to 83.8±4.1 mmHg under global hypoperfusion, with a larger relative change of 9.9±10.4%. But no statistical significance was found (P=0.08). In addition, PO_2_ in the MCA segments was found to be significantly higher than that in the pial collaterals under the control conditions (P=0.023). Shown in Fig. 3C, venous PO_2_ in the MCA territory slightly increased from 42.7±5.4 mmHg under the control conditions to 43.0±4.5 mmHg under global hypoperfusion, with a small relative change of 2.2±19.0%. The watershed counterpart exhibited a larger but insignificant PO_2_ decrease, i.e., from 41.7±3.4 mmHg under the control conditions to 38.2±3.6 mmHg under global hypoperfusion, with a relative change of 8.2±8.7%. Next, PO_2_ was converted to SO_2_, and the oxygen extraction fraction (OEF) was calculated. Fig. 3D shows that the difference in OEF between the control conditions (i.e., 39.8±7.3%) and global hypoperfusion (i.e., 39.7±7.3%) in the MCA territory is negligible. The relative change was calculated as 3.5±29.8%. Strikingly, OEF in the watershed area increased significantly from 36.7±7.6% under the control conditions to 47.4±6.3% under global hypoperfusion, with a much larger relative change of 31.4±16.4% (P=0.006).

Besides, as shown in Fig. 3E, the difference in capillary PO_2_ between the control conditions (48.6±1.9 mmHg) and global hypoperfusion (49.6±2.2 mmHg) in the MCA territory is subtle. The relative change was calculated as 2.1±3.0%. In comparison, capillary PO_2_ in the watershed area noticeably increased from 45.0±3.2 mmHg under the control conditions to 48.6±4.4 mmHg under global hypoperfusion, with a larger relative increase of 8.0±7.1%, but the difference did not reach statistical significance (P=0.058).

### Pial collaterals were dilated under global hypoperfusion

Last, morphological properties of the microvasculature were extracted from the Alexa680-labelled angiograms. Fig. 4B shows that diameter of the pial arteries in the MCA territory was slightly increased from 22.7±2.7 μm under control condition to 23.3±3.0 μm under global hypoperfusion, with a relative change of 2.5±2.4%. Interestingly, the pial collaterals were seen dilated under global hypoperfusion. Qualitatively, diameter of the pial arteries in the watershed area, which are mostly pial collaterals, was significantly increased from 18.9±4.3 μm under control condition to 21.6±3.2 μm under global hypoperfusion (P=0.049), with a relative change of 16.5±12.3%. However, the diameter changes in the pial veins were insignificant (Fig. 4B). And no obvious changes of segment density, diameter and length in either the large vessels (Fig. 4C) or micro-vessels (Fig. 4D) at the deeper layers (100-700 μm) were found.

**Fig. 4.**
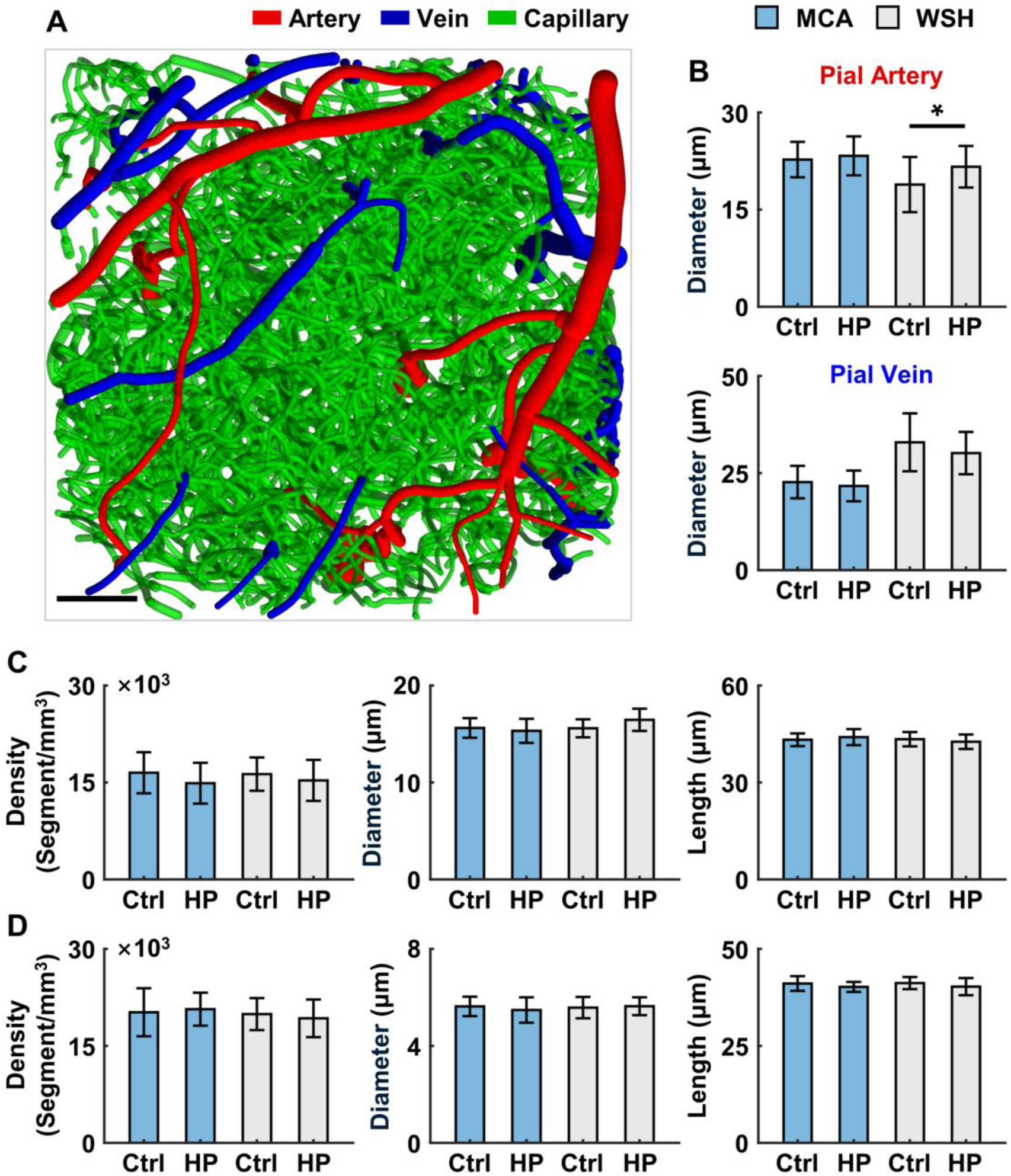
Changes of vascular morphology induced by BCAS. **A.** An anatomical graph showing the top view (X-Y plane) of the microvasculature from the cortical surface down to 600 μm. This graph was generated with the angiogram acquired in the MCA territory, as indicated by the yellow square in Fig. 1A. Scale bar: 100 μm. **B.** Comparisons of vascular diameter. This analysis only included the arteries, veins, and pial collaterals at the cortical surface. **C, D.** Comparisons of vascular segment density, diameter, and length. The analysis included the large vessels with diameter >10 µm (panel **C**) and micro-vessels with diameter ≤10 µm (panel **D**), with the angiograms acquired at the depth range of 100-700 μm under the cortical surface. The data presented in panels **B-D** were acquired in n=4 mice. The same ROIs were chosen for imaging under both conditions. The mean value of each parameter in each ROI (i.e., MCA or WSH) and under each condition (i.e., Ctrl or HP) was calculated first with the angiogram acquired in each mouse, and then across mice. Data are expressed as mean±STD. Asterisk symbol indicates statistical significance (Student’s t-test, P<0.05).

## DISCUSSION

It has been shown that pial collaterals play a significant role in regulating blood reperfusion to alleviate the severity of injury caused by ischemic stroke.^21,42–44^ However, how blood flow and oxygen are distributed in the microvascular networks in the deep cortical layers in the watershed area that is supposedly supported by the pial collaterals has remained elusive. In this work, with multimodal imaging techniques, we quantitatively assessed resting-state blood flow, capillary RBC flux, vascular PO_2_ and morphological characteristics in the cerebral MCA territory and for the first time in the watershed area, in C57BL/6 mice, under normal physiology and chronic global hypoperfusion.

Blood flow in the ascending veins at the cortical surface was measured with Doppler-OCT (Fig. 2A). The presented flow values fall in the range of 0.03-0.1 µL/min, as reported in the previous studies with awake mice.^38,45,46^ Vascular caliber has a strong impact on blood flow. Therefore, it can be reasonably hypothesized that the size of the selected vessels contributes to the flow variation in each imaging region and across regions, as seen in Fig. 2A. Nevertheless, by comparing the flow data acquired in the same vessels under both conditions, we found a BCAS-induced reduction of cerebral blood flow in the two interrogated regions.

Capillary RBC flux was measured by imaging the fluorescence-labelled RBC passages with 2PM (Fig. 2B, C). The mean RBC flux in the gray matter under control condition reported in this work (i.e., ∼67-69 RBC/s in the two interrogated regions; Fig. 2B) is higher compared with the previously published data in awake mice (∼40-50 RBC/s), using the similar imaging approachs.^25,38,45,47,48^ Here, different mouse age, sex, and likely the brain regions chosen for imaging may potentially contribute to the discrepancy. In agreement with our previous studies with both anesthetized and awake mice^31,38^, we observed that RBC flux in the white matter was higher than that in the gray matter under control condition (Fig. 2B, C), especially in the MCA territory where statistical significance was reached (not shown). This could potentially be explained by lower resistance to blood flow in the white matter microvascular networks, leading to higher RBC flux in individual capillary segments. However, blood perfusion in the white matter may still be lower than that in the gray matter, due to much lower capillary density in the white matter.^49–51^ In accordance with venular flow measured at the cortical surface, RBC flux at the deep cortical layers (approximately layers IV and V) as well as in the subcortical white matter was seen reduced under global hypoperfusion in both MCA territory and watershed area. In addition, a significantly larger flux decrease in the white matter was observed (Fig. 2C). These findings are well in line with our previous works, confirming that subcortical white matter is inherently vulnerable to global hypoperfusion.^31,38^ Global hypoperfusion has been regarded as a major cause of white matter diseases and cognitive decline in elders.^52,53^ It has been hypothesized that the vulnerability of white matter to hypoperfusion might be largely due to its distal location along the arterial tree.^31,54,55^ Our results further demonstrate that this effect may be more pronounced in the watershed area, contributing to the development of internal watershed infarcts.^22,56–58^

Furthermore, vascular PO_2_ changes were investigated using 2PM with a novel oxygen-sensitive probe – Oxyphor2P. The baseline PO_2_ data acquired in the arteries and veins at the cortical surface (Fig. 3B, C), as well as in the capillaries at the deeper cortical layers (Fig. 3D) in the MCA territory are all in good agreement with the previous works in C57BL/6 mice of similar age.^25,47^ Our data for the first time show that pial collaterals are, on average, significantly less oxygenated than the neighboring MCA segments under control condition (Fig. 3B). This result can be expected as blood flow in the pial collaterals at rest is relatively slow.^21^ However, the differentially larger PO_2_ changes in both arteries and veins in the watershed area than in the MCA territory ultimately led to a significant OEF increase in the watershed area under global hypoperfusion (Fig. 3D). This observation suggests that a compensatory mechanism was taking place in the watershed area after the BCAS induction, where lower blood flow was compensated by larger oxygen extraction in an attempt to maintain tissue oxygen metabolism and prevent hypoxic injury. The increased OEF and a trend of lower venular PO_2_ imply a decrease in tissue oxygenation in the watershed, likely in the deep cortical layers and subcortical white matter, where the largest decline in capillary RBC flux was observed. However, our intravascular PO_2_ measurements were limited to the upper part of layer V (500 μm), and PO_2_ measurements in the deep cortical tissue are still technically challenging. Therefore, it remains undetermined whether BCAS only resulted in the O_2_ supply-demand mismatch, or it also induced tissue hypoxic islets in the watershed area.

Besides, the morphological parameters (e.g., segment density, diameter, and length) of the microvasculature imaged through the depth range of 100-700 μm are reasonably comparable with the previous studies.^38,59^ However, no significant changes in these parameters have been found. Same as reported in stroke studies^5,21^, the pial collaterals underwent vasodilation during global hypoperfusion (Fig. 4B). This result implies that, potentially, an increased blood pressure gradient along the pial collaterals has been established, facilitating the transport of oxygenated blood from one side to another via the interconnected collateral pathways. The observed trend of increased PO2 in pial collaterals after BCAS procedure (Fig. 3B) is in agreement with potentially increased blood flow in these vascular segments.

This present work has limitations. First, we measured only intravascular PO_2_ in the cortex. Assessments of intravascular PO_2_ in the white matter and brain tissue PO_2_ would be important to determine the appearance of hypoxia and its relation with different microvascular compartments. Second, per the previous studies, BCAS induced a progressive cognitive decline.^28,60^ Thus, it may be important to investigate the evolution of microvascular blood flow and oxygenation over a longer time span. Third, the Doppler-OCT measurements were performed only in venules, as the blood flow calculation in arterioles might be subjected to an aliasing effect due to the high flow speed. Herein, we anticipate that only the measurements of venous blood flow are sufficient to compare the flow changes across different groups, because the total blood flow in arterioles (flowing into the cortical tissue) vs. in venules (flowing out of the cortical tissue) should be comparable, given the size of our interrogated brain area.^61^ Lastly, capillaries were identified empirically based on vascular caliber (≤10 µm) for the measurements of capillary RBC flux and PO_2_. Our fitting criteria with R^2^ ≥ 0.5 ensures the selected vessels follow a single-file flowing pattern, further excluding the smallest arterioles and venules from the analysis.

To summarize, we observed a mismatch between oxygen delivery and consumption in the cerebral watershed area due to global hypoperfusion. We demonstrated that the cerebral watershed area is more vulnerable to global hypoperfusion than the MCA territory, and this vulnerability is greater in the subcortical white matter. The decreased microvascular blood flow under global hypoperfusion in the watershed area is accompanied by a significant increase in OEF, which may potentially contribute to hypoxic brain injury. These changes are associated with the remodeling (e.g., enlargement) of the pial collaterals, which may suggest a novel role of these vessels in mitigating the deleterious effects of global hypoperfusion on the watershed.

## ACKNOWLEDGEMENTS

The authors would like to thank the support from the Ministry of Science and Technology of the People’s Republic of China (2022ZD0211900), the National Institutes of Health (R01NS115401, R01NS121095, R01AG081841, U19NS123717, U24EB028941), and the Science and Technology Innovation Committee of Shenzhen Municipality (JCYJ20220818101611024).

## AUTHOR CONTRIBUTIONS

Baoqiang Li, Sava Sakadzic, and Cenk Ayata designed the study with the help from Ken Arai and Eng Lo; Sava Sakadzic and Baoqiang Li developed the imaging and data-processing methods; Baoqiang Li performed the experiments; Baoqiang Li, Hewei Cao and Yimeng Wu analyzed the data with the guidance from Sava Sakadzic; Baoqiang Li and Sava Sakadzic interpreted the results with the help from Cenk Ayata, Ken Arai, and Eng H. Lo; Srinivasa Rao Allu synthetized the Oxyphor2P probe with the guidance from Sergei A. Vinogradov; Hajime Takase prepared the BCAS model with the guidance from Ken Arai; Buyin Fu performed the animal surgeries for imaging; Baoqiang Li and Sava Sakadzic wrote the manuscript with help from all other authors.

## SUPPLEMENTARY INFORMATION

No supplementary material is provided for this study.

